# Closed-loop fMRI neurofeedback to reduce negative attentional bias in depression

**DOI:** 10.1101/2020.06.07.137943

**Authors:** Anne C. Mennen, Nicholas B. Turk-Browne, Grant Wallace, Darsol Seok, Adna Jaganjac, Janet Stock, Megan T. deBettencourt, Jonathan D. Cohen, Kenneth A. Norman, Yvette I. Sheline

**Affiliations:** Pinceton Neuroscience Institute, Princeton University; Department of Psychology, Yale University; Department of Psychiatry, Perelman School of Medicine, University of Pennsylvania; Institute for Mind and Biology, University of Chicago; Department of Psychology, Princeton University

**Author notes:** Corresponding author: Anne C. Mennen, Princeton Neuroscience Institute, Princeton, NJ 08540.

**Keywords:** real-time fMRI, cognitive training, brain-machine interface, cloud computing, attentional bias, depression

## Abstract

Depressed individuals show an attentional bias toward negatively valenced stimuli and thoughts. Here we present a novel closed-loop neurofeedback procedure that seeks to remediate this bias. Internal attentional states were detected by applying machine learning techniques to fMRI data in real-time, and externalized using a visually presented stimulus that the participant could learn to control. We trained 15 depressed and 12 healthy control participants over three fMRI sessions, preceded and followed by behavioral and clinical assessments. Initially, depressed participants were more likely than non-depressed participants to get “stuck” in negative attentional states, but this diminished with neurofeedback training relative to controls. Depression severity also decreased from pre- to post-training. These results demonstrate that our method is sensitive to the negative attentional bias in depressed individuals, and its reduction after training showcases the potential of this method as a treatment in the future.

## INTRODUCTION

Depressed individuals process negative stimuli differently from healthy participants, leading to differences in attention, memory, and cognitive control (1–5). Depressed participants also tend to show larger and more prolonged neural responses to negative stimuli (6). This may manifest clinically as rumination, or the automatic replay of negative thoughts (1, 7). Given that depressed participants attend more to negative information, researchers have designed paradigms to train participants to reduce this negative bias, and ultimately, depression severity (1, 4).

One common approach is Attention Bias Modification (ABM) training, which involves learning to shift overt spatial attention away from negative stimuli and/or towards positive stimuli (8–12). Another training approach, Cognitive Bias Modification for Interpretation (CBM-I; 13), involves learning to adopt the positive interpretation of an ambiguous situation (e.g., 14).

Following these forms of training, participants typically display the reinforced behavior, for example, attending less to negative stimuli (12, 15). However, transfer to clinical measures (e.g., reduced depression severity or self-reported rumination) has been inconsistent (2, 16–18). A potential limitation of the aforementioned studies is their use of pre-programmed training schedules; recent approaches to attention training have taken a more adaptive approach, providing behavior-based real-time feedback based on mouse position (19) or eye fixation (20–21). Although such approaches have yet to be tested in clinical populations, healthy participants showed promising improvements in reappraisal (21) and rumination (19–20).

While behavioral training has been the main approach to reduce attentional biases in depression, behavioral measures such as button presses and eye movements are downstream effects of underlying neural differences. Neural feedback, such as feedback from functional magnetic resonance imaging (fMRI), allows for measures that are “closer to the source” of the biases, and thus have the potential to be more sensitive and informative. Indeed, depressed participants show neural – but not behavioral – evidence of increased processing of negative stimuli that are presented quickly (see 22). In our work, we therefore sought to combine the advantages of adaptive feedback with the potentially enhanced sensitivity of neural measurements of attention.

A previous study (23) demonstrated the potential of real-time fMRI (rt-fMRI) neurofeedback to improve sustained attention in healthy participants. Participants received visual feedback based on their brain activity during an attention task. Overlaid face and scene images with variable opacity values were shown as participants responded to a cued *go* category and ignored the other, un-cued *no-go* category. Neurofeedback was embedded in a closed-loop circuit: If the neural data indicated that participants were attending more to the incorrect category (e.g., faces), then stimuli in that category were made more opaque (e.g., faces would become more prominent and scenes would become more transparent). This served to “externalize” the participant’s bad attentional state, making the task *more difficult* during attentional lapses, and thus alerting them to try harder to push themselves into a better state. This procedure yielded significant improvements in attention after a single neurofeedback session. Additionally, participants receiving feedback that was veridical (based on their own brain activity) as compared to a control (someone else’s brain activity) exhibited an increased benefit, indicating that individualized feedback was advantageous.

In the current study, we adapted this closed-loop procedure, with the goal of reducing negative attentional bias in depressed participants, rather than improving sustained attention in general. To accomplish this goal, we modified the neurofeedback task so participants always had to ignore negative faces; when participants’ attention drifted to the negative faces, the faces were made more visible and the scenes were made less visible. In this situation, participants need to learn to “unstick” themselves from the negative attentional state in order to make the scenes more visible, so they can continue with the instructed task of responding to the scenes.

Our approach was motivated by a pilot experiment that tested the feasibility of this neurofeedback task in seven depressed participants (24). Based on that work, we refined and expanded the current study to compare two groups (depressed and healthy); we also collected a wider range of behavioral and neural measures to quantify and understand changes over time. Given the technical bottlenecks of real-time fMRI, we implemented the study in open-source Python software for real-time fMRI in the cloud, which we release with this paper to allow other facilities to deploy our method regardless of local computing resources.

We designed our study to (1) establish that our neurofeedback task properly captured negative attentional bias in depression, (2) decrease this bias with training, and (3) show potential transfer to clinical measures. First, we hypothesized that, at the start of training, depressed participants would have difficulty disengaging attention from negative faces. We operationalized this difficulty with a neural measure that tracked the probability of remaining stuck in the most negative attentional state. Second, we hypothesized that neurofeedback training would make it easier for depressed participants to disengage from such negative states. Thus, we expected group differences in sustained negative attention to decrease over time. Third, we hypothesized that this reduction of sustained negative attention would be associated with an improvement in depression symptoms.

## METHODS AND MATERIALS

### Participants

A total of 27 adults participated in the study, including 15 with major depressive disorder (MDD; n female = 8, mean age = 27.3 years) and 12 who served as healthy controls (HC; n female = 6, mean age = 25.4 years). Twenty-three participants completed all 7 visits as planned; two participants could not complete the in-person portions of Visit 6 because data collection was suspended during the COVID pandemic; one participant was lost to follow-up after Visit 5, and one participant was lost to follow-up after Visit 6. For all analyses, we included all data collected, regardless of the availability of follow-up data. Both groups underwent the exact same experimental procedure, differing only in initial diagnosis requirements. Participants were recruited from the University of Pennsylvania Center for Neuromodulation in Depression and Stress (CNDS) laboratory. All participants received monetary compensation for participation. The study was approved by the University of Pennsylvania Institutional Review Board, as well as the Princeton University Institutional Review Board through an IRB Authorization Agreement.

All participants were: 18-60 years old; willing to not take psychotropic medications for the duration of the study; fluent in English; able and willing to provide written consent; and right-handed. Participants in the MDD group scored at least a 16 on the Montgomery-Åsberg Depression Rating Scale (MADRS) clinical interview during Visit 1. Participants in HC group had no history of MDD in their lifetime and no indication of current, significant depressive symptoms (i.e., their MADRS scores were below 8 during Visit 1).

After meeting these eligibility requirements, participants were excluded from both groups if they: had a medical disorder that may cause depression or require medication that could cause depression symptoms, were diagnosed with concurrent DSM-5 psychiatric disorders (e.g., bipolar disorder), had not completed the tenth grade in school, showed active suicide risk, or had MRI contraindications.

### Stimuli

Grayscale face and scene images were adapted from (23). The neutral faces in that study were supplemented with emotional faces from the Chicago Face Database (25), the Karolinska directed emotional faces (26), and NimStim (27); see Supplemental Figure S1 for examples. All faces were cropped to between eyebrows and chin to ensure participants saw emotion and did not rely only on hair while making male/female judgments. Images were then resized to 10° visual angle square and normalized for luminance and contrast. Composite face-scene images with different morphing proportions were generated as in the previous study (23).

### Procedure

The study consisted of seven visits total per participant, as illustrated in Supplemental Table S1. The first five visits were the main study (pre-scanning, 3 neurofeedback sessions, post-scanning). Visit 6 was a behavior-only one-month follow-up. Visit 7 was a three-month follow-up phone call. For all participants, we tried to schedule the first 5 visits as closely as possible within 2 weeks.

Upon providing consent on Visit 1, participants completed the Structured Clinical Interview for DSM-5 Disorders (SCID-5; 28) to assess lifelong symptoms, current depression symptoms, and the presence of additional exclusionary conditions. Participants completed the SCID-5 on Visit 1 only. Upon completion, the MADRS structured clinical interview (29–30) was administered to assess depression symptoms specifically over the week preceding Visit 1. The MADRS was also administered on Visits 5, 6, and 7 to assess how depression severity changed over time. Participants completed additional behavioral and neural tasks before and after neurofeedback (Supplemental Table S1).

Participants completed 7–9 neurofeedback runs per visit on Visits 2-4 (we fit in as many runs as we could within each 2-hour scanning session). Each neurofeedback run contained 8 blocks: the first 4 blocks (“stable blocks”) showed only neutral stimuli with constant opacity and served as training data for the face versus scene classifier; in the last 4 blocks (except run 1), the attended category was neutral scenes and the distractor category was negative faces (Supplemental Figure S1). These blocks served as neurofeedback blocks, in which the opacity changed depending on the relative degree of neural representation of scenes versus faces indicated by a pattern classifier applied to fMRI. Participants were told that the change in opacity was determined by their brain activity rather than their button-pressing accuracy.

At the start of each block, participants were given a cue that indicated the block type (face or scene) and *go* category. For instance, the cue “indoor scenes” indicated that participants should press for indoor scenes (90% *go* trials), and refrain from pressing when seeing outdoor scenes (10% *no-go* trials). Additionally, while making go/no-go judgments, participants had to continuously ignore the overlaid irrelevant stimuli (e.g., faces). For each participant, the *go* scene category (indoor or outdoor) and *go* face category (male or female) were the same across all visits. Assignment of categories was counterbalanced across participants within each group. See Supplemental Figure S1 and (23) for additional task details.

### Data Acquisition

All scanning was acquired with a 3T MRI scanner (Siemens Prisma), using a 64-channel head coil. Sequences were matched to (23) as much as possible. During the first scanning session, we collected a high-resolution magnetization-prepared rapid acquisition gradient-echo (MPRAGE) anatomical scan to construct the whole-brain mask used in real-time and for offline registration. FSL (http://fsl.fmrib.ox.ac.uk/) was used to register the MNI152 standard-space T1-weighted average structure template (31) to each participant’s brain in functional space. We used this registered whole-brain ROI as the mask for each participant. The functional scans consisted of a gradient-echo, echo-planar imaging sequence covering the whole brain (2 s repetition time, 28 ms echo time, 3 mm isotropic voxel size, 64 x 64 matrix, 192 mm field of view, 36 slices). At the end of each scanning session, a fieldmap scan was acquired for offline processing.

### Real-time Processing

Figure 1 provides an overview of our real-time processing system, from the initial stimulus display, to subsequent cloud analysis, and finally, stimulus update. During neurofeedback runs, each new DICOM image was motion corrected to the previous time point following Siemens’ custom motion correction. After that, the DICOM file was saved onto a local Linux machine in the scanning room. Then, the data were masked and flattened into a 1D-vector, and sent to the cloud server for further preprocessing (see Supplement) and classification.

**Figure 1:**
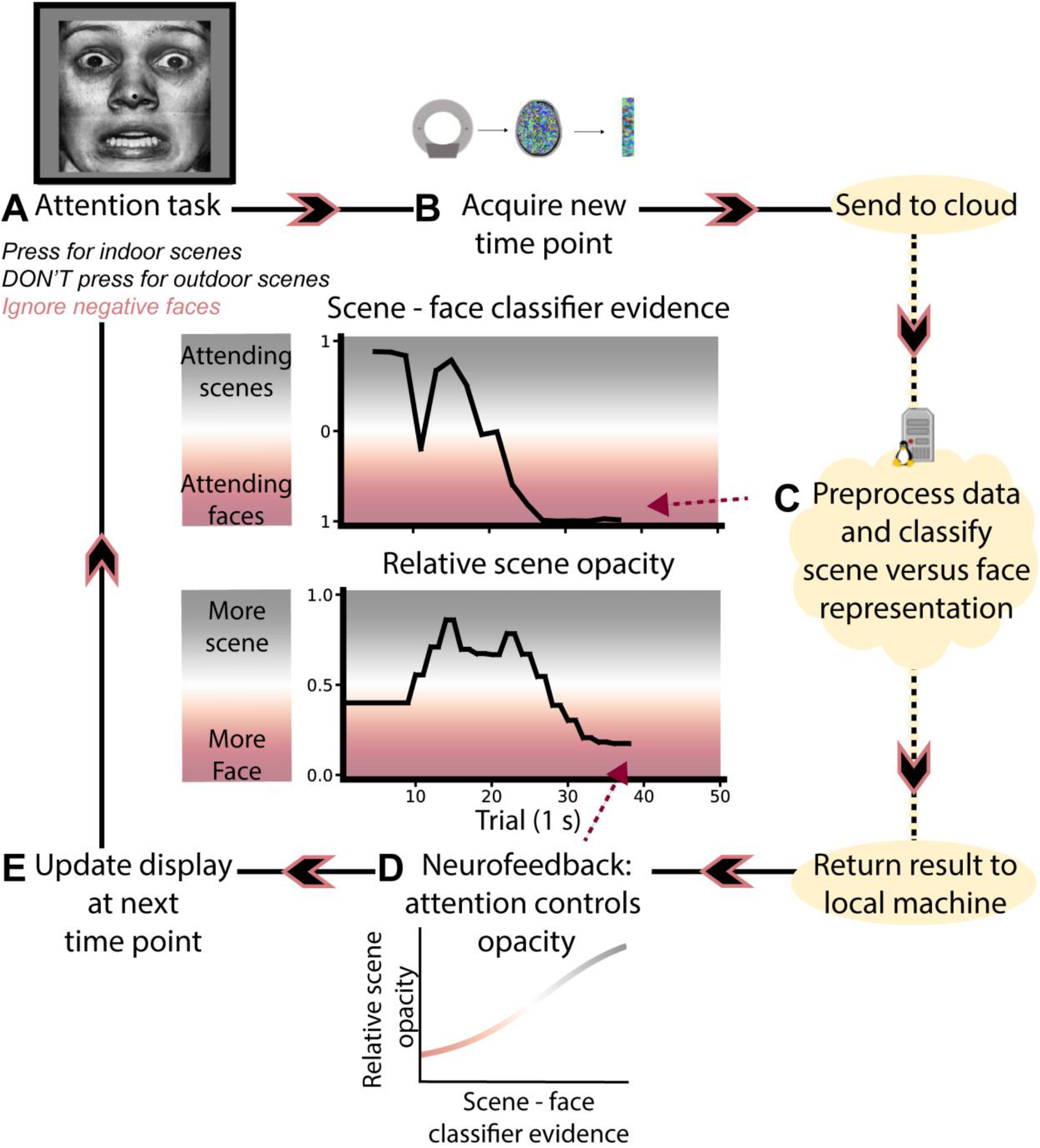
Closed-loop rt-fMRI attention training design. **(A)** Participants perform a go/no-go task on overlaid face/scene stimuli, where they respond based on whether the scene image is indoor or outdoor, and thus have to constantly ignore negative faces. **(B)** As each new time point is acquired, the data are masked and flattened to a 1D-vector. **(C)** The data are sent to a cloud server for preprocessing and classification. **(D)** The result is sent as a text file to the local machine controlling the display. A sigmoidal transfer function converts the relative scene minus face classification evidence difference into opacity proportions, so that the attended category (as measured by the classifier) will become more visually prominent. **(E)** The opacity value is smoothed and updated for the next time point. As shown, when participants are in a maximally negative state, the negative faces dominate the composite image.

During neurofeedback, a multi-voxel pattern classifier (32) was used to decode the extent to which attention was directed at the task-relevant scene or the task-irrelevant face for every time point of image acquisition (TR). The classifier model was re-trained for each neurofeedback run based on the most recent eight stable blocks from before the current run. For example, neurofeedback run 4 used a model trained on the 4 stable blocks from runs 2 and 3 (run 2 used training data from all 8 stable blocks of run 1). Scikit-learn’s logistic regression function (33) was used for model training (with the parameters selected to best match Matlab processing: solver=’saga’, penalty=’l2’, max_iter=300). During neurofeedback, the difference between the amount of classifier evidence for scenes (ranging from 0 to 1) and faces (ranging from 0 to 1) was used as the output neurofeedback score. This score was saved as a text file and sent back to the local computer to influence the display during the following time point.

### Neurofeedback Display

The Matlab script controlling the display loaded each new text file as it was detected. The neurofeedback score was converted to an opacity value for the neutral scene using a sigmoidal transfer function (Figure 1). Then, opacity was smoothed using a moving window over the values from the previous 2 time points to ensure that changes in opacity were not abrupt (23). This smoothed value was set as the opacity for the following 3 trials (1.5 TRs) while the next time point was collected and preprocessed.

### Neurofeedback Performance

To estimate each participant’s attentional state at a given time, we discretized the continuous distribution of scene minus face classifier evidence [ranging from −1 to +1]. Because we used a logistic regression classifier, classification values were distributed toward the extremes (±1). We adjusted the scene minus face classification bins to roughly equate the number of classification samples in each (Figure 2A).

**Figure 2:**
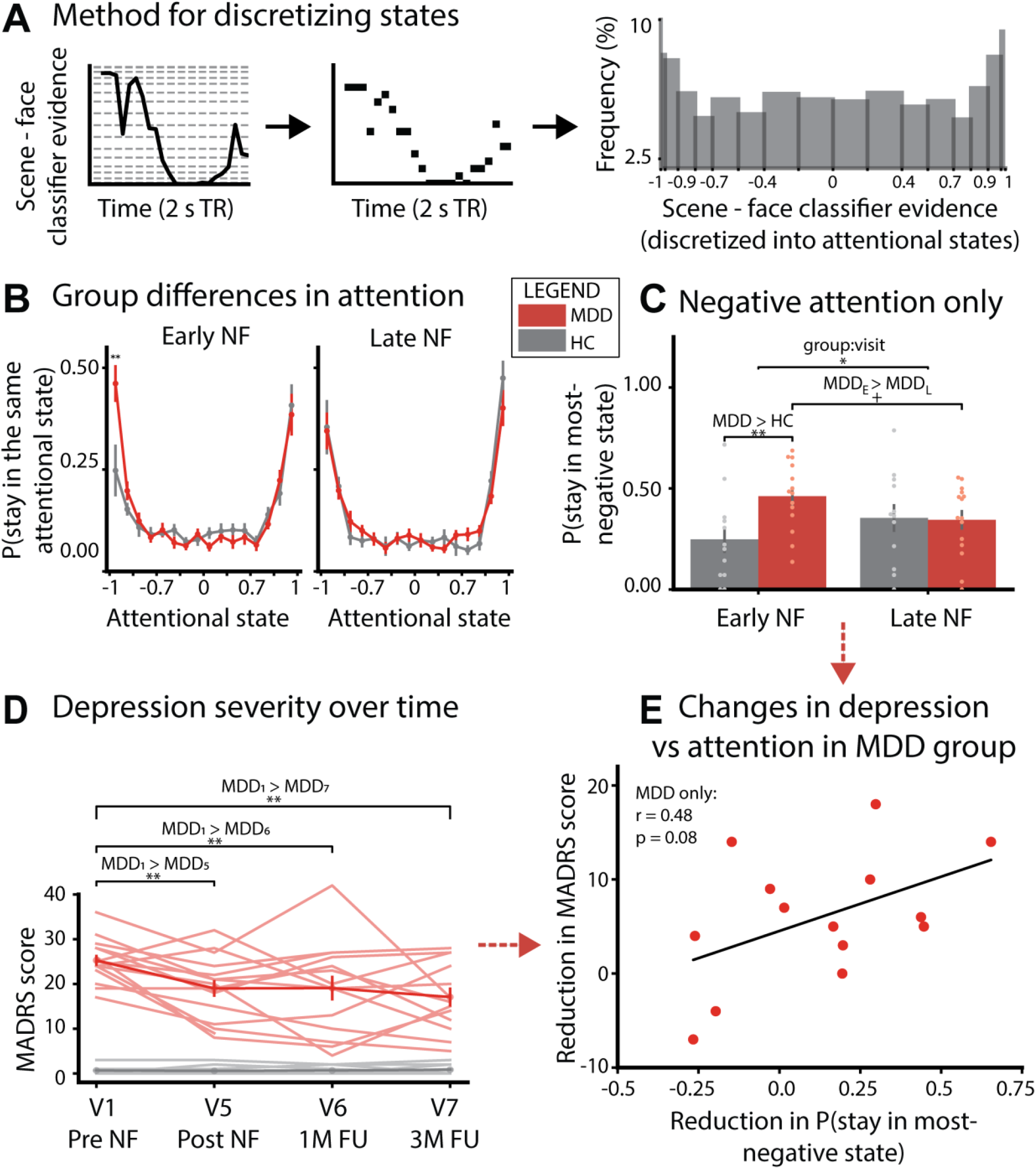
Neurofeedback analysis and results. **(A)** Each block of continuous scene minus face classification evidence was converted into discrete attentional states (dashed lines). This resulted in a roughly equal number of observations in each state across participants. **(B)** Probability of staying in a particular attentional state over time, for Early and Late NF runs. During Early NF, the two groups differed in their probability of staying in the most negative attentional state, plotted separately in **(C)**. This group difference was eliminated by the Late NF runs. **(D)** Depression severity scores significantly decreased for MDD participants over time. **(E)** Within the MDD group, this reduction in getting stuck in the most negative attentional state showed a trending positive correlation with the reduction in depression severity. Circles represent individual participants; bars represent group averages. Error bars represent ±1 s.e.m. ** = p < 0.01; * = p < 0.05; + = p < 0.1

We quantified the extent to which attentional states persisted over time. We operationalized this measure as the conditional probability that the scene minus face difference remained in the same bin across time points. Because we were interested in how feedback delivered at time *t* affected attention, we compared the attentional state at time *t* to the attentional state at time *t* plus 5 seconds (where we would expect feedback effects on brain activity to be maximal, accounting for the hemodynamic lag). As each TR was 2 s, we separately calculated results using 2- and 3-TR shifts, and averaged the results to estimate a 5-s shift. Specifically, for each time delay, *d* (2 or 3 TRs), for a given attentional state bin, *A*, we calculated the persistence of that state by counting the number of times that the scene minus face classification value fell within *A* at time *t + d*, given it was in *A* at time *t*. We then divided this number by the total number of occurrences of state *A*, shown in the equation below:

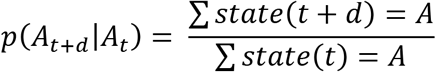

We then averaged all conditional probabilities from the 2 and 3 TR shifts over all runs considered. To understand how attention changed from early to late in training, we isolated the initial three neurofeedback runs of the first session (Visit 2; Early NF) and the final three neurofeedback runs of the last session (Visit 4; Late NF).

## RESULTS

### Depression severity

As hypothesized, depression severity decreased over time for MDD participants (Figure 2D). Depression scores decreased significantly from pre-training in Visit 1 to post-training in Visit 5 (one-tailed t(14) = 3.61, p = 0.0014), to the one-month follow-up in Visit 6 (one-tailed t(13) = 2.85, p = 0.0069), and to the three-month follow-up in Visit 7 (one-tailed t(12) = 3.43, p = 0.0025).

At the start of neurofeedback training, the largest difference between groups in the probability of remaining in the same attentional state over time occurred for the most negative state, with MDD participants showing a greater tendency to get stuck in this state (one-tailed t(24) = 2.80, p = 0.0049)^1^ (Figure 2B).

At the end of neurofeedback training, MDD participants were marginally less likely to get stuck in the most negative state, compared to the start of neurofeedback training (paired one-tailed t(13) = 1.67, p = 0.059). Additionally, there was a significant interaction between group and visit, such that the MDD group decreased in their probability of getting stuck in the most negative state from early to late in training relative to the HC group (unpaired one-tailed t-test comparing change in MDD group to change in HC group, t(24) = 2.04, p = 0.026; Figure 2C).

Additionally, the reduction in the MDD group was associated (across participants) with a marginally significant reduction in depression severity (Pearson r = 0.48, p = 0.083) (Figure 2E). There was also a trending correlation in the same direction in the HC group (Pearson r = 0.51; p = 0.090).

## DISCUSSION

Our closed-loop neurofeedback method successfully detected the difficulty that depressed participants have in disengaging attention from negative stimuli – this was evident in our finding that, at the outset of training, MDD (vs. HC) participants were more likely to get “stuck” in the most negative attentional state (Figures 2B, 2C). Of note, there were no significant initial differences in the *average* level of scene minus face classifier evidence for MDD vs. HC participants (Supplemental Figure S2) – that is, it was not the case that MDD participants simply had more brain activity related to negative faces. To expose the initial difference between groups, we had to rely on a measure that specifically tracked participants’ tendency to persist in a negative state.

It is also noteworthy that there were no initial differences in *behavioral* performance on the go/no-go task between MDD and HC participants (Supplemental Figure S3). The detection of group differences for the go/no-go task in neural – but not behavioral – data implies that neural measures may be more sensitive for capturing attentional differences. This underscores the value of using rt-fMRI neurofeedback training to reduce negative attentional bias.

Compared to most depression studies in the rt-fMRI literature, this study is unique in its design and analysis methods. First, we trained participants to disengage attention from negative stimuli; by contrast, most of the previous rt-fMRI studies trained depressed participants to increase neural responses to *happy* stimuli, such as images (34–35) or autobiographical memories (36–39). Regulating positive emotions has yielded robust benefits. For example, (35) even found unintentional clinical benefits for the control group, who imagined relaxing scenes while regulating scene-specific ROIs. Our study explored a less-common approach of training away from negative stimuli, instead of toward positive stimuli. This was based on our belief that learning to regulate negative attention may strike at the underlying dysfunction more directly. The relative efficacy of training negative vs. positive attention can be tested in future studies, e.g., by using a variant of our paradigm where participants are instructed to attend to positively valenced faces or scenes, while ignoring neutral distractors from the other category.

Our study differs from the small number of other rt-fMRI studies using negative stimuli in that we regulated a decoded cognitive state, as opposed to mean ROI activity (40–41). For instance, (40) trained 10 depressed participants to reduce neural responses to negative images within an individualized ROI (defined based on the single voxel most sensitive to negative images within the “salience network”). In another study, (41) had participants recall negative memories while using a strategy from cognitive behavioral therapy (CBT). During CBT application, a neurofeedback signal was used to train participants to decrease anterior cingulate cortex activity. Both studies yielded promising clinical benefits specific to real-time training, in the form of decreased negative self-descriptions (40) and increased use of the trained CBT strategy after neurofeedback (41). More work is needed to assess the relative efficacy of our closed-loop attention-training procedure compared to ROI-based approaches.

Another avenue for future work is to verify that the positive clinical effects we observed were specific to the individualized nature of the neurofeedback. To demonstrate this, one could recruit a follow-up control group of MDD participants. These participants would receive feedback scores that are either yoked to a previous depressed participant’s brain (23) or determined by an irrelevant ROI (36–37). If this group does not show the same improvements, we can be more certain that the improvements shown by our MDD participants relate to receiving individualized neurofeedback (vs. a more general effect of the procedure).

An important feature of the method reported here is that the real-time analysis of the imaging data performed on the cloud. Although the sophistication and practical utility of real-time fMRI has increased over the past 10 years (42), the prevalence of its use has been hampered by hardware requirements and by the technical complexity of setting up and running an experiment. Offloading computation to the cloud should help to make this approach accessible to researchers regardless of local computing resources and local computational expertise. In future work, we plan to extend our framework to control both the real-time processing and neurofeedback display via a cloud-based web server, to further minimize local dependency.

In summary, this study highlights the potential clinical benefits of real-time fMRI neurofeedback procedures that target specific cognitive states. By tracking sustained attention over time, our procedure provides a face-valid way of detecting the difficulties that MDD patients experience in getting “stuck” in negative states. This was borne out in the observed sensitivity of our measure to initial differences between MDD and HC participants. By “externalizing” these internal attentional lapses (i.e., making task-irrelevant negative faces more visible as they were attended more), our procedure provides rich feedback that patients can leverage to learn to control these states. This training potential is supported by our findings showing reduced sustained negative attention in MDD patients and reduced depressive symptoms. By making this technique openly accessible on the cloud, we hope to make it easier for other researchers to explore the benefits of this approach in diagnosing and treating MDD and other clinical syndromes.^2^

## ACKNOWLEDGEMENTS

This work was supported by funding from Intel Corporation and the John Templeton Foundation (JDC, KAN and NTB), National Institutes of Health grant T32MH065214 (ACM), the Canadian Institute for Advanced Research (NTB), and the University of Pennsylvania Endowment (YIS). The opinions expressed in this publication do not necessarily reflect the views of these organizations. The funders had no role in study design, data collection and analysis, or decision to publish. We thank Paula Brooks, Mark Elliot, Jordan Gunn, Neggin Keshavarzian, Arlene Lormestoire, Elizabeth McDevitt, Kevin Pendo, and Coco Zhao for assistance in running the study. We thank Mihai Capotă, Kai Li, Daniel Suo, Yida Wang, and Ted Willke for discussions and early contributions to software development and hardware systems for real-time fMRI. We also give special thanks to David Schnyer and Christopher Beevers for their key contributions to the Schnyer et al. (2015) pilot study that launched this line of work.

## Supplemental Information

## INTRODUCTION

As mentioned in the main text, our study included additional neural and behavioral tasks to assess the ability of each task to capture initial group differences and changes in performance over time. Here, we present details regarding our full study design, cloud processing, and results.

## STUDY DESIGN

**Table S1:**
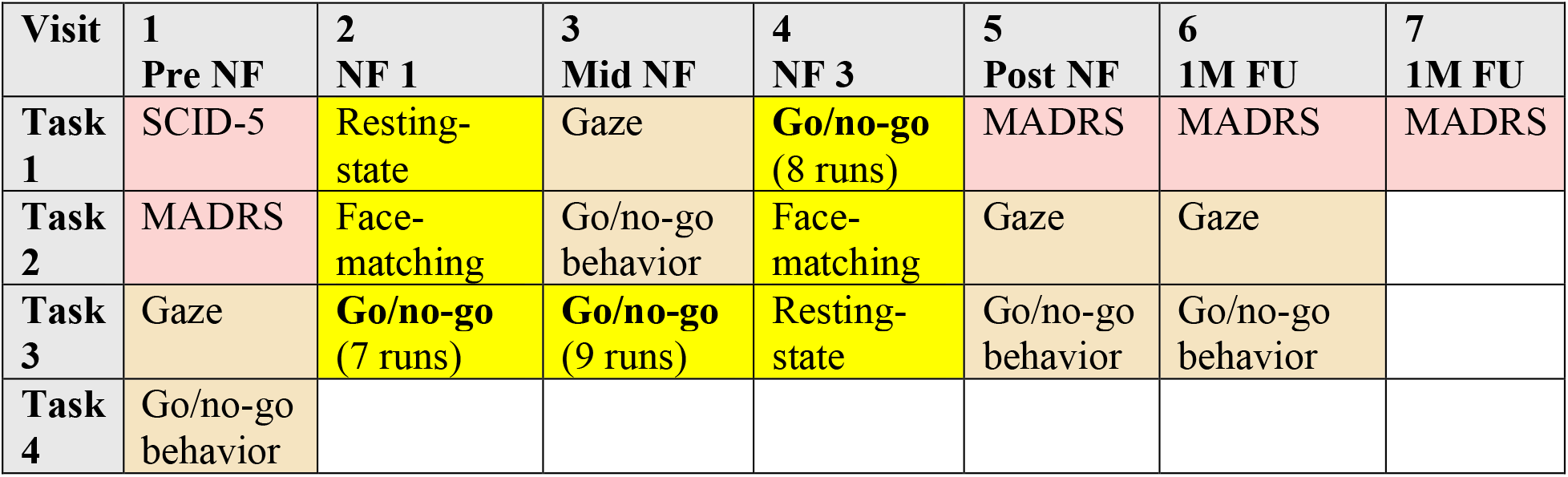
Full study design for all 7 visits: pre-neurofeedback, three neurofeedback sessions, post-neurofeedback, a one-month follow-up, and a three-month follow-up visit. Columns separate the different visits, while rows indicate the order of tasks performed at each visit. Cell color specifies the type of task: **pink** = clinical assessments; **beige** = behavioral tasks completed outside of the scanner; **yellow** = scanner tasks. **NF** = neurofeedback; **FU** = follow-up. Go/no-go in **bold** is added to emphasize the rt-fMRI neurofeedback runs.

| Visit | 1<br>Pre NF | 2<br>NF 1 | 3<br>Mid NF | 4<br>NF 3 | 5<br>Post NF | 6<br>1M FU | 7<br>1M FU |
| --- | --- | --- | --- | --- | --- | --- | --- |
| Task 1 | SCID-5 | Resting-state | Gaze | **Go/no-go**<br>(8 runs) | MADRS | MADRS | MADRS |
| Task 2 | MADRS | Face-matching | Go/no-go behavior | Face-matching | Gaze | Gaze |  |
| Task 3 | Gaze | **Go/no-go**<br>(7 runs) | **Go/no-go**<br>(9 runs) | Resting-state | Go/no-go behavior | Go/no-go behavior |  |
| Task 4 | Go/no-go behavior |  |  |  |  |  |  |

As shown in Table S1, participants completed 7 visits to assess clinical, behavioral, and neural changes over time. For all behavior-only tasks, stimuli were presented on a 60 x 34 cm monitor (1280 x 1024 resolution) with the Psychophysics Toolbox for Matlab (http://psychtoolbox.org/). For all scanning tasks, stimuli were presented on a projector with 1280 x 720 resolution. Of note, we collected resting state scans for a portion of the participants. The first 2 healthy control (HC) participants and the first 5 depressed (MDD) participants did not receive resting state scans. Because we were missing resting state scans for a substantial number of participants, we do not analyze those scans here.

### Go/no-go task design

We used an adapted version of the go/no-go task as described in (1) for behavioral and neural assessments of sustained attention.

**Figure S1:**
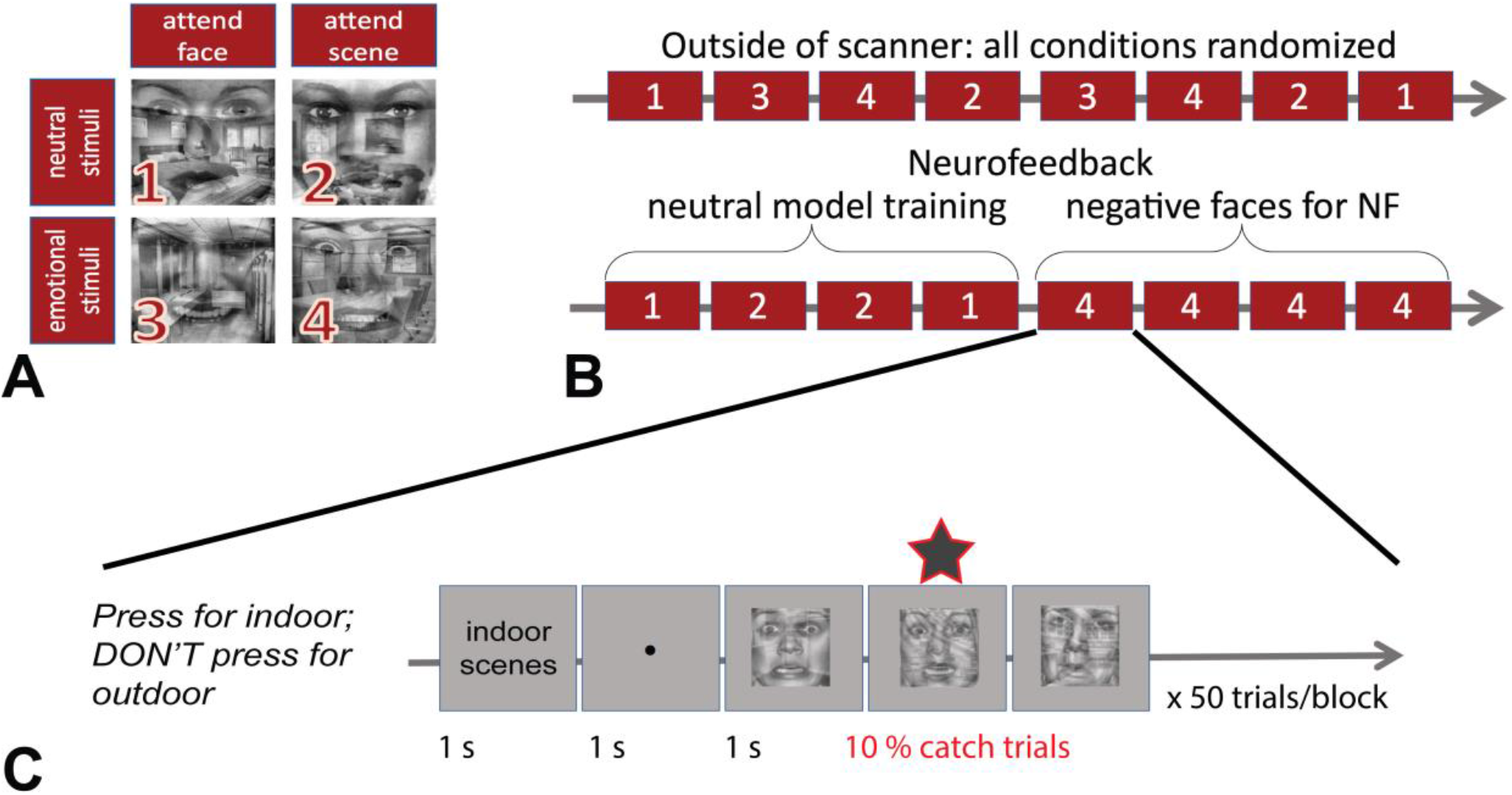
Go/no-go task design and differences in the neurofeedback and behavior-only version. **(A)** In all conditions, participants saw an overlaid face/scene image, while having to attend to one category and ignore the other category. The columns specify which category was attended (face or scene), and the rows specify if the faces were neutral or emotional. Altogether, the 4 conditions were: **(1)** attend neutral faces; ignore neutral scenes, **(2)** attend neutral scenes; ignore neutral faces, **(3)** attend happy faces, ignore neutral scenes, **(4)** attend neutral scenes; ignore negative faces. **(B)** Participants completed 8 blocks of the task during each run. In the behavior-only version, all conditions were presented in a randomized order. In the neurofeedback version, participants first saw neutral faces and scenes. During neurofeedback, they only saw negative faces and neutral scenes. The version with happy faces and neutral scenes could be used in the future to assess the relative benefit of training attention toward positive stimuli versus away from negative stimuli. **(C)** Example trial structure during the go/no-go task.

## CLOUD PROCESSING

Cloud processing was developed and implemented so that the analysis of DICOM images was no longer reliant upon local resources or proprietary software. We executed the Python-based code on a Microsoft Azure virtual environment that was part of Penn Medicine’s secure network.

Cloud processing involved sending the masked 1D-vector of each volume of DICOM data to a virtual machine. Once the data arrived, images were spatially smoothed with a Gaussian kernel FWHM of 5 mm. During the last 4 blocks of each run (the neurofeedback blocks), each incoming 1D-vector was high-pass filtered in real-time (using all time points in the current run up to that point; cutoff = 200 s), and z-scored using the mean and standard deviation from the previous 4 stable blocks of the current run, as in (1).

*NOTE*: Before cloud processing was completed, all real-time preprocessing and classification were performed identically to (2). This was the case for HC participants 1-2 and MDD participants 1-6, who received Matlab-based neurofeedback. All other participants received feedback from Python-based cloud processing, unless a specific technological issue (e.g., the internet being down) obstructed us from using the cloud. In that case, Matlab was used. To ensure that the processing method did not bias results, for all neurofeedback analyses, we went back and processed all data with the same real-time cloud pipeline. The mean Pearson correlation between the results obtained with this uniform pipeline and the results that were actually used for neurofeedback was 0.996, 95% CI [0.995, 0.997].

## ADDITIONAL NEUROFEEDBACK RESULTS

### Average classification

In addition to computing attentional states, we also looked for differences in average scene minus face classifier evidence (Figure S2). We did not observe classification differences between MDD and HC participants, either at the beginning of neurofeedback (t(25) = 0.32, p = 0.76) or at the end of neurofeedback (t(25) = 0.34, p = 0.74). On average, both groups showed a decrease in average scene minus face classification from early to late neurofeedback (combined paired t(26) = 2.61, p = 0.015).

**Figure S2:**
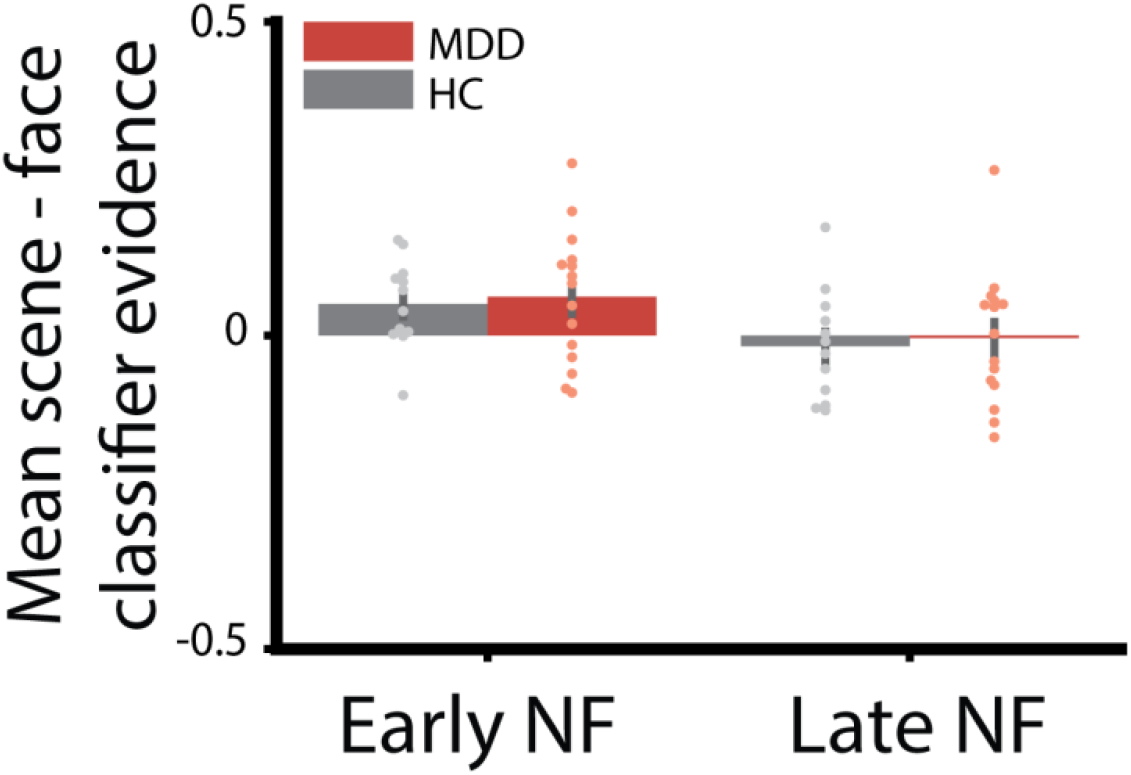
Average scene - face classifier evidence during early and late neurofeedback runs. No group differences were found. Circles represent individual participants; bars represent group averages. Error bars represent ±1 s.e.m.

## GO/NO-GO BEHAVIOR TASK

### Background

The go/no-go behavior task was used to measure attentional biases through behavioral key presses. Again, the task consisted of overlaid face and scene images, though this time with constant, equal opacity. As with the neurofeedback task, participants always were instructed to attend one category (e.g., neutral faces) and ignore the opposite category (e.g., neutral scenes). For this behavioral version, we also included a condition where participants attended to happy faces and ignored neutral scenes (Figure S1).

To quantify sustained attention through behavior, we used the same measure of sensitivity (A’; 3) that we used in (1). In our study, we calculated A’ differences between different emotional conditions (e.g., ignoring neutral faces versus ignoring negative faces). Before neurofeedback, we expected that MDD participants would have a more difficult time, and thus worse A’ performance, when ignoring negative faces compared to neutral faces.

We also hypothesized that MDD participants might show more difficulty (i.e., lower A’ scores) when attending to happy faces compared to attending to neutral faces. While there is some evidence to suggest that depressed participants show deficits attending to positive stimuli, attention to positive versus negative stimuli is usually compared directly (4–6). For this reason, it is unclear if the difference in attention is due to a lack of attention to positive stimuli or a bias to attend to negative stimuli.

### Methods

Participants completed 4 runs of this version of the go/no-go task during Visits 1, 3, 5, and 6. Each run contained 8 50-s blocks. As shown in Figure S1A, there were 4 stimulus conditions in total. Thus, each of the 4 conditions was shown twice during each 8-block run. The condition order was randomized for each participant within each run (Figure S1B).

To measure sustained attention with behavior, we estimated performance according to the nonparametric measure A’ (1). This measure tracks sensitivity to the *go* versus *no-go* categories, factoring in both hit and false alarm rates. To estimate differences in performance when participants have to ignore negative faces compared to neutral faces, we calculated the negative bias as

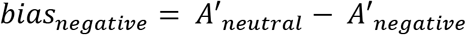

Thus, negative biases greater than 0 indicate *worse* performance when ignoring negative faces, compared to neutral faces. This would suggest problems with disengaging attention away from negative stimuli.

Likewise, we estimated differences in performance when participants have to attend to positive faces compared to neutral faces as

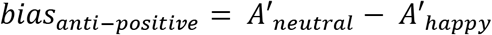

Thus, anti-positive biases greater than 0 indicate *worse* performance when attending to happy faces, compared to neutral faces, which would suggest difficulty in focusing attention on happy stimuli.

### Results

**Figure S3:**
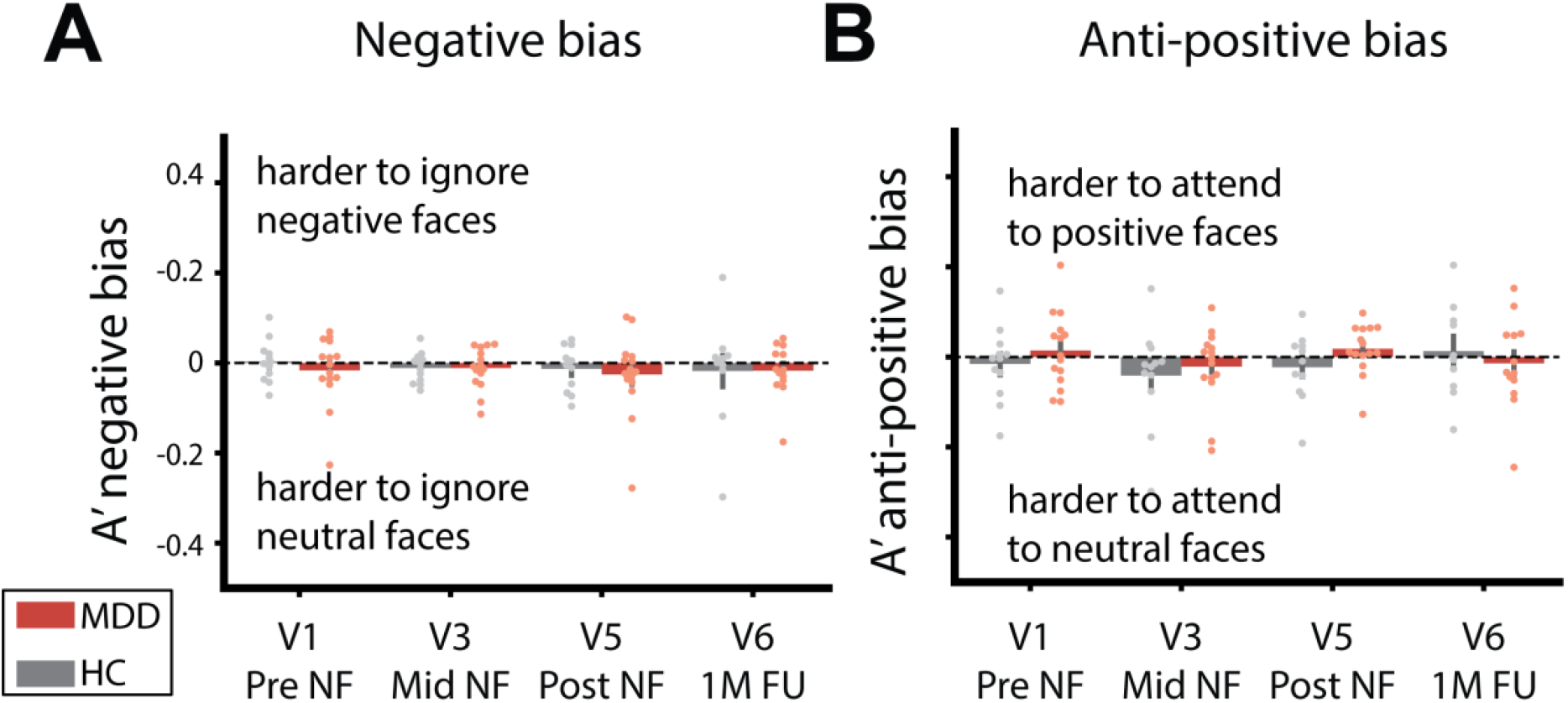
Results for the go/no-go behavior task for **(A)** negative and **(B)** anti-positive images. No significant group differences were evident for either attentional bias metric at any visit. Circles represent individual participants; bars represent group averages. Error bars represent ±1 s.e.m.

As shown in Figure S3A, we found no group differences during Visit 1 (t(25) = 0.78, p = 0.44), nor at any point during the course of training for the A’ negative bias. We did not find any group differences in A’ anti-positive bias during Visit 1 (t(25) = −0.94, p = 0.36), nor at any stage during training (Figure S3B).

## GAZE TASK

### Background

This task was designed to measure sustained attention with eye-tracking data, which allows for a continuous estimate of attention. Based on (6), each trial displayed 4 images representing different emotions (dysphoric, threatening, happy, and neutral) on the screen at once for 30 s. Recording eye movements throughout these long trials allowed us to assess which stages of attention (e.g., orientation, maintenance) differed the most between groups before training, and if those group differences changed over time.

Before neurofeedback, we expected MDD participants to show difficulty disengaging attention away from the dysphoric images (7–8). This would manifest as prolonged attention to the dysphoric images, as was the case in (6) and (4). Given initial group differences, we expected the negative bias to decrease over time, demonstrating that the benefits of rt-fMRI neurofeedback transferred to this task.

### Methods

First, we adapted (6) for a multi-session design by creating 4 different versions of the task to be completed on Visits 1, 3, 5, and 6 (Table S1). We supplemented images from the original study with stimuli from the International Affective Picture System (IAPS; 9) and the Nencki Affective Picture System (10). We included all IAPS images used in the original version (6).

To normalize images across databases, the images were cropped to be square. Next, images were sorted into four different versions while balancing for image entropy. We then made sure that all versions of the task did not differ significantly in entropy, contrast, or luminance. All participants completed each version, but the version order was randomized for each participant.

A Tobii X120 eye-tracker (Tobii Technology, Stockholm, Sweden) was used to record eye movements (sampling rate = 120 Hz). Participants’ chins were fixed to a free-standing chin rest during the task to ensure that they had limited head motion and maintained a constant distance from the monitor. After eye-tracking calibration, participants were instructed to freely view the different images presented on the screen as if they were watching television or looking at pictures in a photo album (as in 6). We told participants that we were tracking their eyes to make sure they were attending to the images on the screen.

As in (6), the task consisted of 20 trials, with each trial lasting 30 s. Each trial was either a neutral filler or a valence trial. During neutral filler trials, the 4 images were all neutral. During valence trials, there was a neutral, dysphoric, threatening, and happy image shown on the screen. The full task consisted of 8 neutral filler trials and 12 valence trials that were presented in a random order for each participant. As mentioned above, participants completed 4 different versions of this task. See Figure S4 for trial structure and example stimuli.

*Note*: Various obstacles (eye makeup, pupil size, glasses, etc.) impeded continuous eye position estimates for some participants. We adapted to the difference in measurements by normalizing our estimates by the number of gaze fixations that were recorded, as described next.

**Figure S4:**
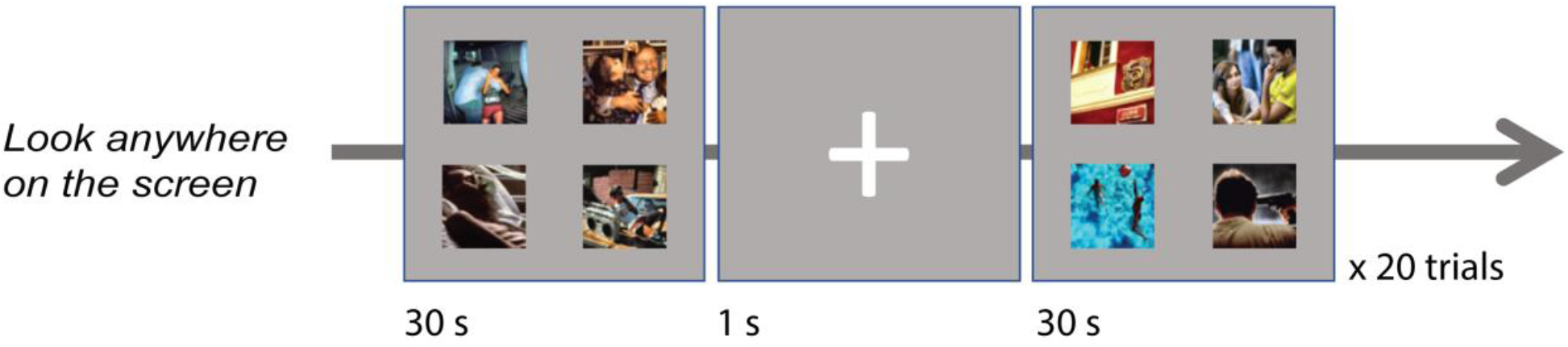
Trial structure for the gaze task. Participants were instructed to freely view the images as if they were watching television. During each trial, 4 images were shown on the screen at once. Eye movements during the full 30-s trial were recorded to estimate sustained attention over time. Note: both of the example trials shown are valence trials, as there is a dysphoric, threatening, happy, and neutral image on the screen. Neutral filler trials were the same, except that all 4 images were neutral. The task consisted of 20 trials total: 12 valence trials and 8 neutral filler trials.

We recorded the (X,Y) positions from each eye during the 30-s trials. Following (6), we preprocessed the eye-tracking data by defining eye positions in terms of *fixations*. Specifically, we filtered the data such that fixations had to (1) be maintained for at least 100 ms and (2) remain within 1° of visual angle for this duration. After filtering all samples into a time series of fixations that met this criteria, each fixation was classified according to the attended emotional image. Because there were unequal numbers of fixations for a given participant across trials, we normalized each of the metrics described next by the number of data points collected.

## INITIAL ORIENTATION

To assess initial orientation differences by image category and group, for each trial, we recorded the image category upon which the participant first fixated. We calculated the probability of initial fixation for each day by taking the total number of times that the participant first fixated on that image category, divided by the total number of times that the participant fixated on any of the images. This way, we did not count the trials in which no viable data were collected.

## TOTAL VIEWING RATIO

To calculate the total viewing ratio of dysphoric images, for each trial, we totaled the number of fixations on dysphoric images divided by the total number of fixations recorded for that trial. We then averaged across all trials for each day.

## MAINTENANCE VIEWING RATIO

To understand if participants had difficulty *disengaging* attention away from dysphoric stimuli, for each trial, we counted the number of fixations on the dysphoric image, given that the participant was already fixated on the image. If the participant fixated on the dysphoric image, viewed other images, and then returned to the dysphoric image, we only considered the first continuous fixation in our analysis. We then divided this number of samples by the number of fixations recorded for that trial. Again, this normalization step corrected for the lack of data in some trials for some participants. Finally, we averaged the maintenance ratios across all trials to get one ratio per day per participant.

### Results

**Figure S5:**
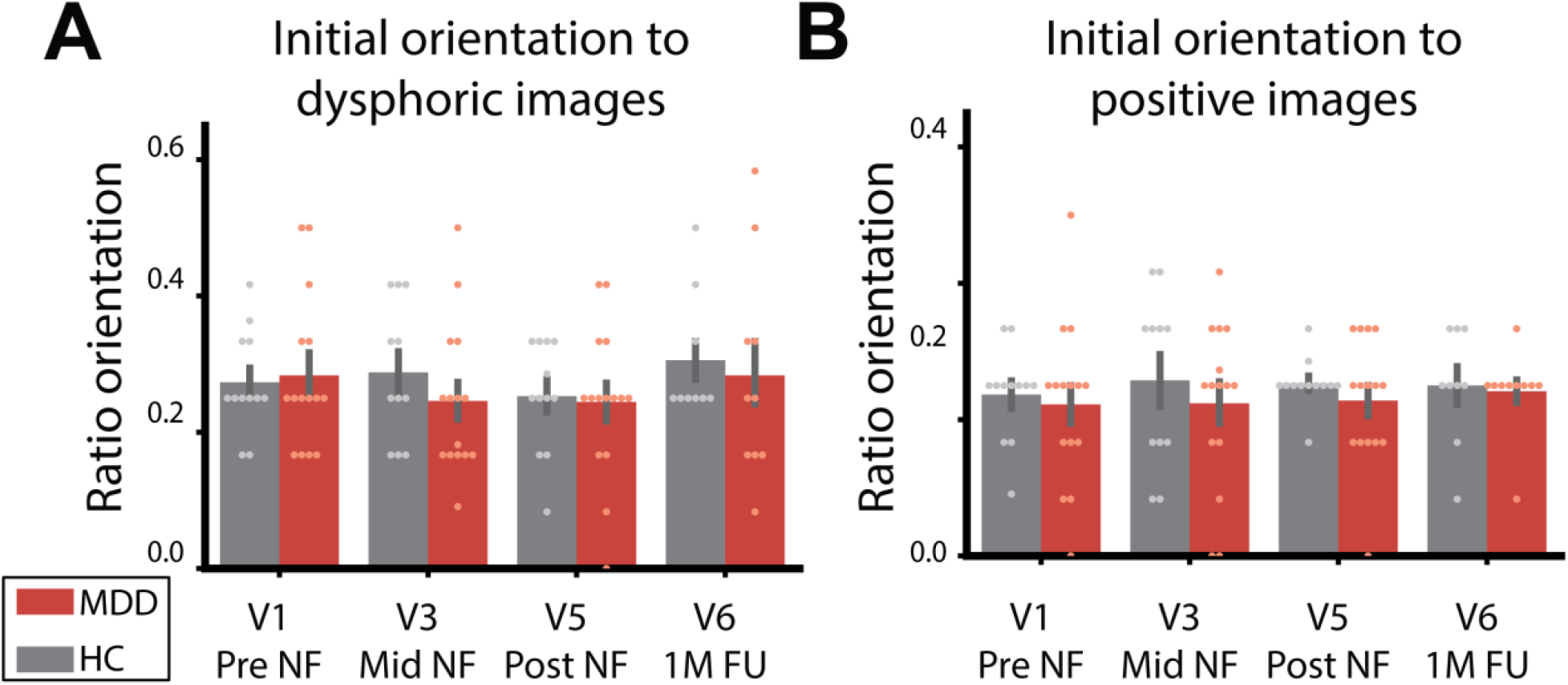
Initial orientation to **(A)** dysphoric and **(B)** positive images. No significant differences were found between groups at the onset of training, or on subsequent visits. Circles represent individual participants; bars represent group averages. Error bars represent ±1 s.e.m.

## INITIAL ORIENTATION

We did not find group differences in the initial orientation measure. On Visit 1 (Pre NF), MDD participants did not orient to dysphoric images significantly more often (one-tailed t(25) = 0.26, p = 0.40; Figure S5A), nor did MDD participants orient to positive images significantly less often (one-tailed t(25) = −0.37, p = 0.36; Figure S5B). There were no significant differences between groups on subsequent visits either.

**Figure S6:**
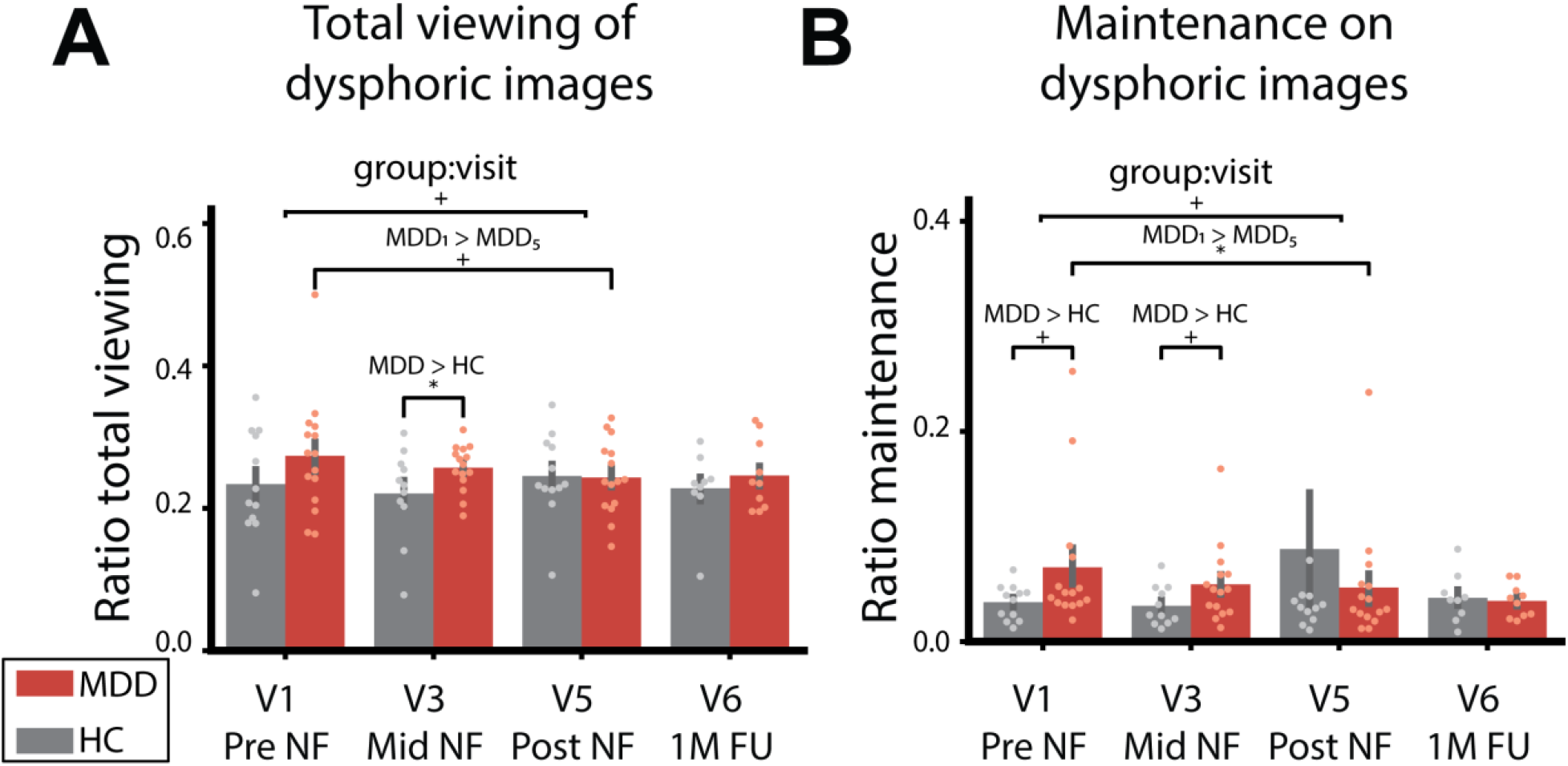
**(A)** Total viewing ratio for dysphoric images **(B)** Maintenance ratio on dysphoric images. MDD participants reduced both metrics on average from pre- to post-neurofeedback, but initial group differences and MDD improvements were larger for maintenance on dysphoric images. Note: One outlier in the HC group only fixated once during V5 and thus that data point is not shown within the axis limits in (B). Circles represent individual participants; bars represent group averages. Error bars represent ±1 s.e.m. * = p < 0.05; + = p < 0.1

## TOTAL VIEWING RATIO

On Visit 1, MDD participants spent more total time viewing dysphoric images compared to HC participants on average, though this was not significant (one-tailed t(25) = 1.28, p = 0.11). Additionally, MDD participants showed a trending decrease in the total amount of time they spent viewing dysphoric images from pre- to post-neurofeedback (paired one-tailed t(14) = 1.44, p = 0.086). Considering changes between Visits 1 and 5, we found a trending but not significant interaction between group and visit (unpaired one-tailed t-test comparing change in MDD group to change in HC group, t(25) = 1.33, p = 0.098). Figure S6A plots group performance for all visits.

Because MDD participants decreased in the amount of the time spent viewing dysphoric images, we examined the relationship between the decrease in total time spent viewing dysphoric images and the decrease in MADRS scores. There was a slight positive relationship in the MDD participants but it was far from significant (Pearson r = 0.30, p = 0.28).

## MAINTENANCE VIEWING RATIO

On Visit 1, there was a marginally significant effect where MDD participants maintained fixation for a longer amount of time compared to HC participants (one-tailed t(25) = 1.69, p = 0.051). From pre- to post-neurofeedback, MDD participants significantly decreased their maintenance time on dysphoric images (paired one-tailed t(14) = 1.85, p = 0.043). Considering changes between Visits 1 and 5 only, we found a trending but not significant interaction between group and visit (unpaired one-tailed t-test comparing change in MDD group to change in HC group, t(25) = 1.44, p = 0.082). Figure S6B shows group performance over all visits.

As the MDD participants decreased their maintenance time on dysphoric images and also showed decreased depression severity after neurofeedback, we examined the relationship between these improvements in the MDD participants. There was no significant relationship between these measures (Pearson r = −0.033, p = 0.91).

## FACE-MATCHING TASK

### Background

We included the face-matching task to assess group differences in neural responses to emotional faces. This task was adapted from (11) to measure neural responses to negative, neutral, and happy faces in a block design. Originally, (11) analyzed group differences in *average* amygdala responses to negative relative to neutral blocks. Before neurofeedback, we expected MDD participants to show higher limbic system reactivity during negative face blocks (11–13).

However, because we were interested in sustained attention, we were unsure if averaged block activity would capture negative biases related to disengagement. We referred to (14) for an additional analysis approach, which found that depressed participants showed increased amygdala responses to negative images *only* towards the end of the trial. This motivated us to consider the time course of amygdala activity, as an increased response towards the end of the block may indicate a failure to inhibit processing of negative stimuli. Given initial group differences in the amygdala response either over the whole block or at the end of block only, we expected the group differences to decrease after neurofeedback through transfer learning.

### Methods

**Figure S7:**
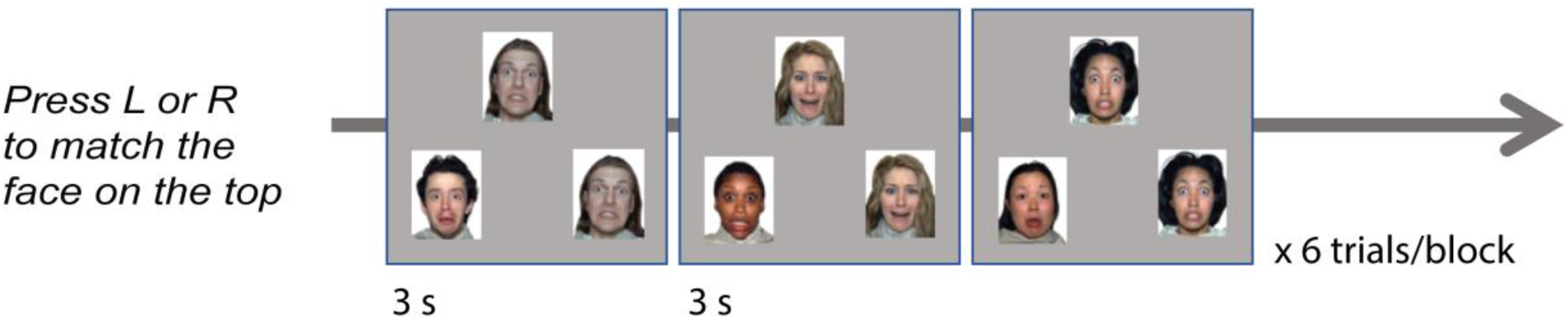
Trial structure for the face-matching task. Participants pressed their index or middle fingers to match the top image to the left or right bottom image, respectively. There were 6 3-s trials within each block. All block types (happy faces, fearful faces, neutral faces, objects, and fixation) repeated 3 times within a run. Shown above is an example of a fearful face block.

We included the same face and object images as in (11) Face stimuli were adapted from the Radboud (15) and NimStim (16) stimulus sets. There were 2 versions of this task; one version drew images from the Radboud set and the other used images from the NimStim set. As participants completed this task twice (Visits 2 and 4; Table S1), the order in which each participant completed the tasks was randomized by participant.

PsychoPy2 (17) was used for this task. Participants completed 2 5-minute runs of this task during Visits 2 and 4. The task involved matching images in 18-s blocks. There were 5 block types of different stimulus categories: happy faces, fearful faces, neutral faces, objects, and fixation. Each block type repeated 3 times within a run. Participants were instructed to choose the bottom image that matched the top image by pressing their index fingers to indicate a left match or their middle fingers to indicate a right match (11). Figure S7 shows the trial structure in each block.

After scanning, data were stored following the Brain Imaging Data Structure (BIDS) specification for further processing (18). Then, fMRIPrep (19) was used for subsequent offline processing and image registration.

Results included in this section come from preprocessing performed using fMRIPprep 1.0.11 (RRID:SCR_016216; 20-21), which is based on Nipype 1.1.6-dev (RRID:SCR_002502, 22-23). The following description of preprocessing was generated with fMRIPrep.

### Anatomical data preprocessing

Each T1w (T1-weighted) volume was corrected for INU (intensity non-uniformity) using N4BiasFieldCorrection (ANTs 2.1.0; 24) and skull-stripped using antsBrainExtraction.sh (ANTs 2.1.0), using the OASIS template. Brain surfaces were reconstructed using recon-all from FreeSurfer 6.0.0 (RRID:SCR_001847; 25), and the brain mask estimated previously was refined with a custom variation of the method to reconcile ANTs-derived and FreeSurfer-derived segmentations of the cortical gray-matter of Mindboggle (RRID:SCR_002438; 26). Spatial normalization to the ICBM 152 Nonlinear Asymmetrical template version 2009c (RRID:SCR_008796; 27) was performed through nonlinear registration with the antsRegistration tool of ANTs 2.1.0 (RRID:SCR_004757; 28), using brain-extracted versions of both T1w volume and template. Brain tissue segmentation of cerebrospinal fluid (CSF), white-matter (WM) and gray-matter (GM) was performed on the brain-extracted T1w using fast (FSL 5.0.9, RRID:SCR_002823; 29).

### Functional data processing

Functional data were slice time corrected using 3dTshift from AFNI (AFNI 16.2.07; 30) and motion corrected using mcflirt (FSL 5.0.9; 31). Distortion correction was performed using fieldmaps processed with fugue (FSL 5.0.9; 32). When there was no fieldmap scan due to ending a scan short (for one participant on one session), we used “fieldmap-less” distortion correction. This was performed by co-registering the functional image to the same-participant T1w image with intensity inverted (33–34) constrained with an average fieldmap template (35), implemented with antsRegistration (ANTs).

After distortion correction, this was followed by co-registration to the corresponding T1w using boundary-based registration with 9 degrees of freedom, using bbregister (Freesurfer 6.0.0; 36). Motion correcting transformations, field distortion correcting warp, BOLD-to-T1w transformation and T1w-to-template MNI warp were concatenated and applied in a single step using antsApplyTransforms (ANTs 2.1.0) using Lanczos interpolation (37). Framewise displacement (38) was calculated for each functional run using the implementation of Nipype.

Many internal operations of fMRIPrep use Nilearn 0.4.2 (RRID:SCR_001362; 39), principally within the BOLD-processing workflow. For more details of the pipeline, see the <u>section corresponding to workflows in fMRIPrep’s documentation</u>.

After data were preprocessed and transformed into MNI space with fMRIprep, we extracted confounds for removal. For each run of the task, we extracted the 6 motion parameters along with framewise displacement. Processing included (1) removing the first 5 TRs of each run, (2) spatially smoothing with (FWHM = 6mm) using nilearn (39), and (3) z-scoring each voxel over time. The two runs completed on each day were combined for further processing with 3dTproject in AFNI (40) which involved: high-pass filtering (cutoff = 100 s) and regressing out motion confounds. Although calculating beta coefficients for block types as in (11) would be an appropriate way to analyze the data, we did not calculate beta coefficients so that we could extract responses over time. We sought to replicate the findings in (14) specifically that amygdala activity continued to rise for MDD participants towards the end of viewing sad faces.

Because we were interested in limbic response to negative faces, we focused on the left amygdala (LA). To derive the LA ROI we repeated the following steps for each participant: (1) we registered all aparc+aseg labels to functional T1w space using Freesurfer’s mri_convert and mri_label2vol functions (25), (2) we extracted the LA from the individual Freesurfer labels in native space, (3) as a final step, we converted the amygdala mask to MNI space using the ANTs-derived registration matrix from fMRIPrep (28). Finally, we created an aggregate LA mask by intersecting all of the amygdala masks from each participant.

To compare amygdala activity across groups, we averaged the time series from each block within the LA ROI. We then subtracted the resulting time series between the negative and neutral blocks. To consider the entire hemodynamic response, we included 5 TRs before the block start and 5 TRs after the block start.

### Results

As described above, we expected MDD participants to show increased attention to negative faces, resulting in larger and more prolonged responses in the LA ROI (14). If we found this initial difference in processing negative images, and our neurofeedback manipulation successfully reduced negative attentional bias, we would expect the difference between groups to decrease after neurofeedback.

First, we looked to see if average activity over the entire block differed by group before neurofeedback. We shifted the average amygdala signal by 2 TRs (4 seconds) to account for the HRF delay. Then, we averaged activity over the entire block (18 s, or 9 TRs). On average, MDD participants did not show larger activity to negative versus neutral faces before neurofeedback (one-tailed t(25) = 0.96, p = 0.17).

Next, instead of considering the average activity over an entire block, we evaluated if the time series showed group differences. Figure S8A plots the average time series for each group for negative – neutral blocks before neurofeedback. As shown, the response to negative faces in the MDD group increases over time. Eventually, the group difference grows to a significant level one TR (unshifted) into the next block (one-tailed t(25) = 1.95, p = 0.032).

As shown in Figure S8B, after neurofeedback, MDD participants did not show an increased LA response for negative faces one TR into the next block (one-tailed t(25) = 0.35, p = 0.36). From pre- to post-neurofeedback, MDD participants showed a trending decrease in their response at this TR (paired one-tailed t(14) = 1.57, p = 0.069), but HC participants did not (paired one-tailed t(11) = −0.034, p = 0.49). We did not find a significant interaction between group and visit for this specific TR response (unpaired one-tailed t-test comparing change in MDD group to change in HC group, t(25) = 1.074, p = 0.15).

Even though MDD participants decreased their LA response to negative faces from before to after neurofeedback, we did not find a significant relationship between decreases in LA activity and reduction in MADRS scores (Pearson r = −0.047, p = 0.87).

**Figure S8:**
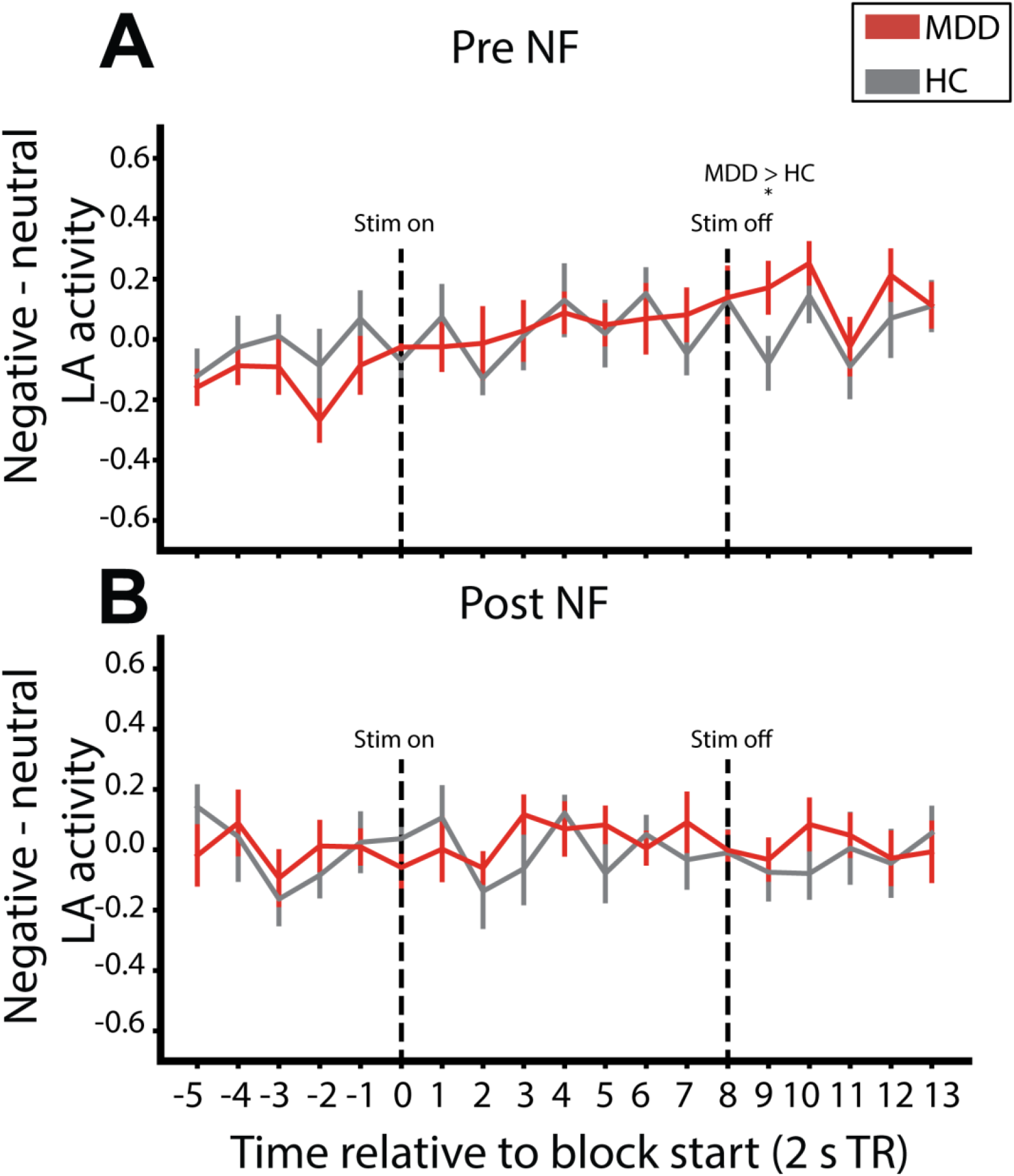
Left amygdala response to negative - neutral face blocks. **(A)** Pre NF responses **(B)** Post NF responses. MDD participants showed increased LA signal during negative face blocks before neurofeedback, but not after. Lines represent group averages. Error bars represent ±1 s.e.m. * = p < 0.05

## Footnotes

1 One of the MDD participants was never in the most negative attentional state during the Early NF period, so the P(stay in most-negative state) measure was undefined for this participant during Early NF; consequently, this participant was omitted from analyses involving the Early NF period.

2 Links to software and documentation: (1) display code (https://github.com/amennen/rtAttenPenn_display), (2) rt-cloud processing code (https://github.com/brainiak/rtAttenPenn_cloud), (3) documentation for running an experiment (https://docs.google.com/document/d/1mI9S-5GYjOfDwFT5Ewb7nWyHPoE00GsI0K0uk0_l6UA/edit?usp=sharing), and (4) example DICOM data (https://doi.org/10.5281/zenodo.3873446)

